# Glycocalyx dynamics and membrane curvature under cytosolic pressure

**DOI:** 10.1101/2023.05.14.540721

**Authors:** Jay G. Gandhi, Donald L. Koch, Matthew J. Paszek

## Abstract

The glycocalyx is a soft material composed of glycosylated proteins and lipids decorating the plasma membrane of cells. Although the glycocalyx is a mechanical conduit between the plasma membrane and the cell surroundings, the coupled dynamics of the glycocalyx and membrane are poorly understood in most cell processes. Here, we construct a dynamic model to predict the coupled mechanical behaviors of the glycocalyx and membrane due to internal cytosolic pressure in a cell interacting with an external substrate. We report how the glycocalyx constituents bear cytosolic loads, physically rearrange, and shape the cell membrane. Simulations of the model predict that highly flexible, polymeric elements in the glycocalyx are uniformly compressed in the cell-substrate contact zone. However, more rigid, fiber-like glycocalyx constituents dynamically rearrange over timescales on the order of 10 milliseconds, concentrating into clusters. This spatial heterogeneity allows close apposition of the plasma membrane with the substrate, a geometry that would be favorable for subsequent adhesive bond formation. Clustered glycocalyx constituents imprint nanotopographical features on the plasma membrane with longitudinal dimensions of 200 – 300 nm. Analysis of the membrane topographies reveals curvatures that could be sufficient to elicit biological responses through curvature sensing proteins, such as BAR-domain proteins. Together, our simulations suggest how instabilities in the compressed glycocalyx could mediate downstream adhesive and signaling processes in the bleb-substrate interface.

## INTRODUCTION

The plasma membrane of most eukaryotic cells is covered in a dense meshwork of proteins and glycans that comprise the cellular glycocalyx. The glycocalyx commonly includes glycosaminoglycans (GAGs) and large polymeric glycoproteins called mucins, which have been implicated in various biological contexts. For instance, cancer aggression is correlated with the increased expression of several cell-surface mucins, including Muc1, Muc16, and podocalyxin, as well as some GAGs (1–3). Mucins are also commonly expressed on the surface of multiple immune cell types with amoeboid migratory phenotypes, including activated T-cells and dendritic cells (4–6).

The glycocalyx has been known to act as a physical barrier between cells and extracellular components such as the extra-cellular matrix (ECM) and other cells (3). The glycocalyx biopolymer Muc1 is known to guard the cell against attack from immune cells (7), where the thickness and the strength of the protective barrier is modulated by the density and the length of the glycocalyx polymers (7). The Muc1 glycopolymer has also been shown to regulate the formation of focal adhesion complexes and subsequent biochemical signaling pathways by modulating the physical distance between the cell membrane and extracellular substrates (2). Macromolecular crowding outside the cell surface is also known to increase retention time scales for soluble factors in extracellular media and enhance their reaction rates (8).

Embedded in the plasma membrane, the glycocalyx has a close relationship with the shape and curvature of the cell membrane as well as processes impacted by membrane curvature. The crowding of bulky glycocalyx polymers spontaneously generates curvature in the plasma membrane, leading to membrane shapes such as blebs, tubes, and pearls depending on the packing density of the glycocalyx biopolymers (9). Mechanically loaded GAGs and mucins can also imprint topographical features on the plasma membrane (2). Membrane topography is associated with spatiotemporal regulation of adhesive and signaling processes. Distances between the membrane and the substrate dictate the kinetics of adhesive bond formation, with the kinetics of interactions slowing exponentially with increased separation between receptors and ligands (2). Membrane curvatures can stimulate the reorganization of lipids, lipidated proteins and transmembrane proteins, as well as regulate protein function (10). Imprinting membrane topography through compression against nanoscale extracellular features can recruit BAR-domain containing proteins to activate curvature sensing pathways, resulting in local induction of actin polymerization and associated cellular responses (11). Furthermore, local membrane invaginations can enhance rates of clathrin-mediated endocytosis (12). While theoretical descriptions of membrane bending are relatively abundant in the literature (13–15), incorporation of these alongside a modeling framework for the glycocalyx has been explored only recently.

Cells *in vivo* frequently exhibit fast, amoeboid migration phenotypes using spherical leading-edge protrusions called blebs (16,17). Embryonic, immune, and metastatic tumor cells commonly adopt a blebbing migratory phenotype for efficient locomotion in three-dimensional (3D) extracellular matrices and tissues (1,18). The creation of such blebs is driven by actomyosin contraction and the resulting hydrostatic pressure on the plasma membrane (19). The plasma membrane peels away from the underlying cytoskeleton, allowing the membrane to rapidly expand into a spherical projection (20). Blebs can grow to sizes of up to approximately 2 μm in ∼30 seconds in response to internal bleb pressures of up to 100 Pa (21,22). GAGs and cell-surface mucins are directly implicated in the induction of blebs and other protrusions of the plasma membrane, suggesting that these macromolecules may be present on blebs of diverse cellular origin (9). High molecular weight GAGs and mucins are typically the largest elements in the glycocalyx and, thus, would form the initial physical contacts between a protruding bleb and an extracellular substrate. As a bleb contacts the extracellular substrate during blebbing motility, these macromolecules would be mechanically loaded by the cytoplasmic pressures that drive bleb expansion (23). The dynamic behavior of the glycocalyx is particularly understudied in the context of these sustained cytoplasmic pressures. Moreover, the role of the glycocalyx in determining the curvature of an expanding bleb has not been investigated.

Despite the central role of the glycocalyx in mediating interactions between cells and their surroundings, insufficient analyses exist that elucidate the physical mechanisms at play in the glycocalyx during such interactions. In particular, the role of the glycocalyx in governing the cell membrane topography under the presence of cytosolic pressures is unknown. In this manuscript, we simulate the constituents of the glycocalyx and the cell membrane under the presence of a cytosolic pressure. The simulations in this paper may offer insights into the roles of the glycocalyx during biomechanical processes such as membrane curvature generation and bleb extension.

The work here builds upon the modeling framework presented in (23) and predicts the deformation and organization of the glycocalyx as well as the membrane shape. We consider the glycocalyx to be either an array of beams or a brush of flexible chains to cover the broad range of molecular stiffnesses observed for GAGs and mucins. The glycocalyx elements can move laterally in the plane of the membrane and rearrange due to compressive stresses. A shape equation captures bending deformations of the cell membrane. We consider the dynamic transport of the glycocalyx constituents due to convective and diffusive fluxes, including osmotic pressures in the glycocalyx. The next section provides a summary of the modeling framework used in this paper.

## MODEL

### Summary

In this manuscript, we construct a framework for simulating the interplay between the glycocalyx and cell membrane in the presence of a cytosolic pressure (Fig. 1). The modeling framework for the dynamic simulations in this paper builds on an earlier model for equilibrium simulations of the glycocalyx (23). Cell-surface mucins and GAGs are idealized as polyelectrolytes and comprise the mechanical elements in the glycocalyx. We consider two limiting treatments for the polyelectrolytes as either flexible chains in a polymer brush or a continuum of bendable beams. We consider each of these cases at once to understand the physical behavior of the glycocalyx at the two limits of polymer flexibility. Physical behavior of a mixture of the beam glycocalyx and the brush glycocalyx is not captured here and would require separate treatment. We account for membrane bending using beam bending theory. We incorporate steric effects between the polyelectrolytes in the glycocalyx as well as the osmotic pressures of bulk counterions neutralizing the charged residues in the glycocalyx. The growth of a cytosol-filled bleb interfacing with an extracellular matrix is a dynamic process having the potential for important transient processes and kinetically trapped static states. Therefore, in this paper we carry out dynamic simulations of the glycocalyx. Out of the possible transient processes during the cytosolic compression of the glycocalyx, the lateral transport of the glycocalyx polyelectrolytes is the process with the limiting time scale. Therefore, we consider an unsteady transport equation that describes the spatiotemporal dependence of the glycocalyx density. The next few sections contain descriptions of the modeling elements that combine to describe the dynamics of a cytosolically compressed glycocalyx.

**Fig. 1.**
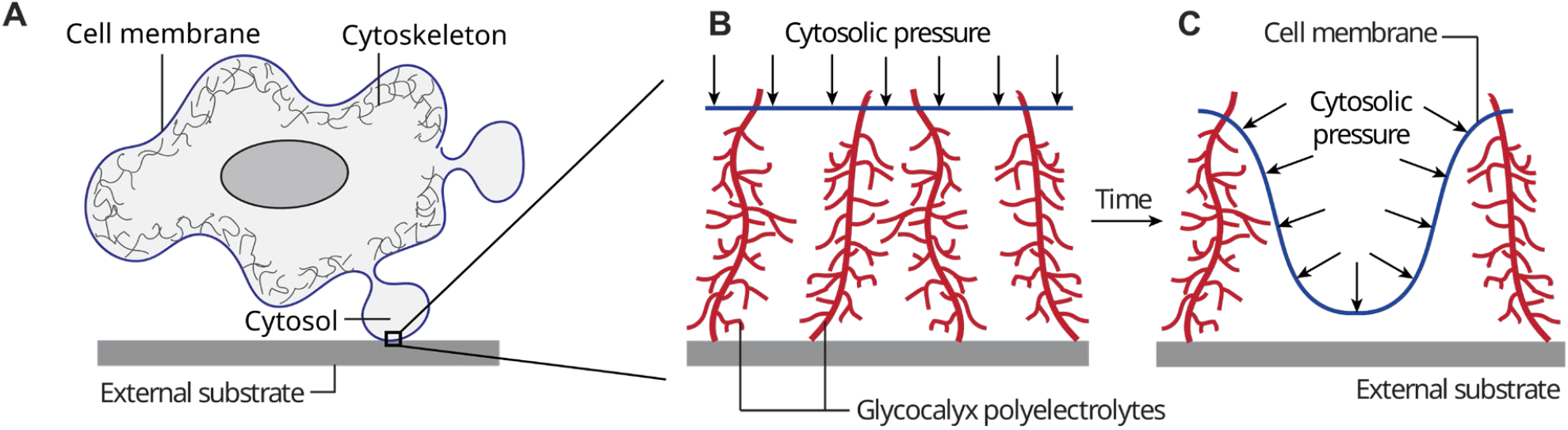
A schematic showing expected physical behavior of the glycocalyx on a cell membrane filled with cytosolic fluid and interacting with an external substrate. **A**. A cell exhibiting cytosol-filled protrusions interacting with an extracellular substrate. The small black box shows a nanoscale region on the cell interacting with the substrate. **B**. In the nanoscale region, the glycocalyx and the cell membrane act as the interface between the cell and the substrate. Thus, the glycocalyx and the cell membrane experience experience the cytosolic pressure from the cell. **C**. Given enough time, the cytosolic pressure is predicted to push the glycocalyx out and create membrane locations with close proximity to the external substrate as well as membrane regions with high curvature.

### Cell membrane mechanics with cytosolic pressure

The membrane is deformed due to the forces acting on the membrane, namely the cytosolic pressure, reaction forces from the glycocalyx, force dipoles exerted by the glycocalyx, and the osmotic pressure on the membrane. We consider the cell membrane during the growth of a bleb, during which the cytoskeleton is absent from the bleb membrane. Hence, we assume cytoskeletal forces to be absent on the membrane. The shape of the cell membrane can be captured by the Helfrich equation (13,14,24,25). Balancing the forces and torques driving membrane bending with the internal resistance of the membrane to bending results in the membrane shape equation. We consider a form that allows for large bending deformations of the membrane and we employ the constant surface area assumption typically used in membrane bending models (13,24–26). Cell membranes are known to offer strong resistance to lateral stretching or compression and instead prefer undergoing bending deformations (27). We consider the membrane to offer a linearly elastic response to bending deformations (13–15). Thus, the constitutive relation for the membrane is 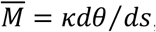, where 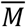 is the internal bending moment generated by the membrane, *κ* is the bending modulus of the membrane, *θ* is the local inclination of the surface from the horizontal substrate, *s* is an arc-length coordinate on the membrane. *dθ*/ *ds* is the local membrane curvature. Fig. 2A shows a generic configuration of the membrane considered in the modeling framework. Note that although Fig. 2B-C show a truncated diagram of the membrane, the entire membrane domain is simulated. We consider a portion of the membrane domain that is much smaller than the bleb size. Hence, we consider the membrane to be flat at the onset of the cytosolic pressure.

**Fig. 2.**
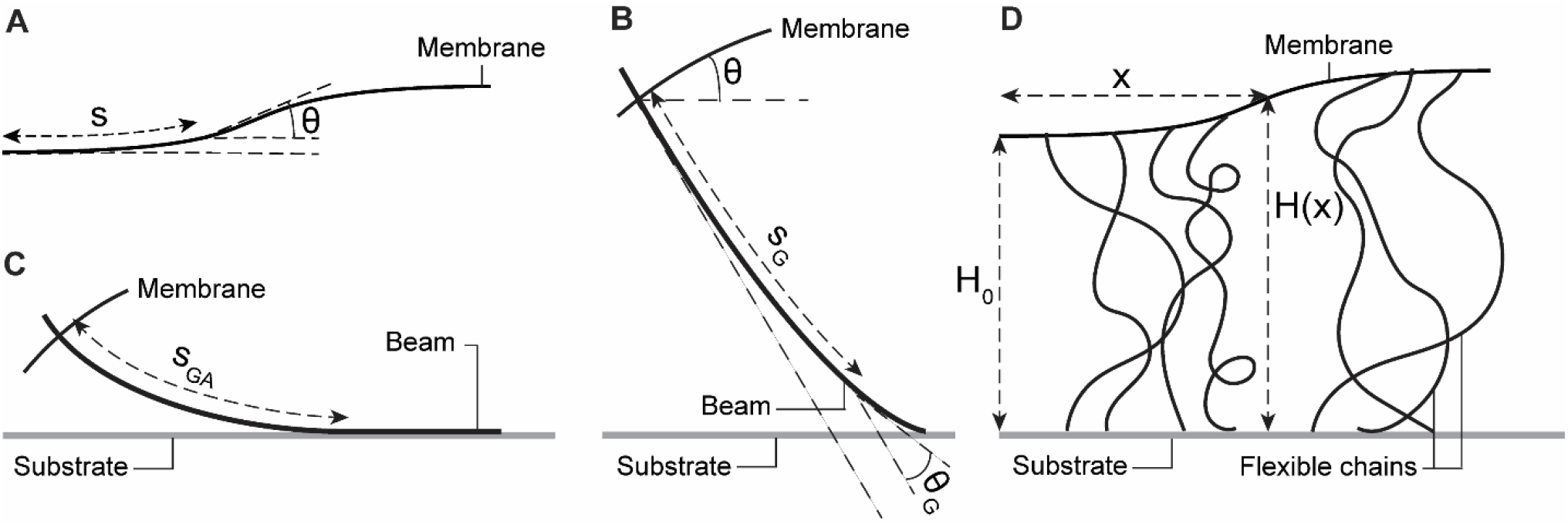
Schematics of physical modeling components. These schematics depict the mechanics-derived models used to describe the shape of the cell membrane and the beam-like and chain-like structural elements of the glycocalyx. **A**. Arc length coordinates, *θ*(*s*), are used to describe the shape of the membrane domain. **B**. A separate arc length coordinate system, *θ*_*G*_ (*s*_*G*_), is used to describe the shape of each beam-like glycocalyx polyelectrolyte. Note that **B** shows only a truncated piece of the cell membrane in order to focus attention on the glycocalyx polyelectrolyte. **C**. As a special case of **B**, for glycoprotein forces above a transition load, the part of the beam-like glycoprotein beyond the arc length *s*_*GA*_ aligns with the flat external substrate. **C** also shows only a truncated diagram of the membrane. The entire domain of the membrane shown in **A** is considered in the simulations. **D**. Flexible chains in the glycocalyx can adopt a distribution of entropy-governed configurations and are modeled using the polymer brush framework.

The cytosolic pressure acts over the length scale of the membrane domain and is locally normal to the membrane. We include a term, *P*_*I*_, designating the intracellular pressure relative to a uniform pressure outside the cell. The cytosolic pressure, *P*_*I*_, appears in the membrane equation alongside the other normal forces, *P*_*N*_, on the membrane. *P*_*N*_ is the net pressure experienced by the membrane due to counterion and polymer brush osmotic pressures. *P*_*N*_ is locally normal to the membrane. The pressure difference between the ends of the membrane domain is expected to be much smaller than the pressure difference across the membrane. Hence, we consider the cytosolic pressure, *P*_*I*_, to be spatially uniform. The moment balance on the membrane is:

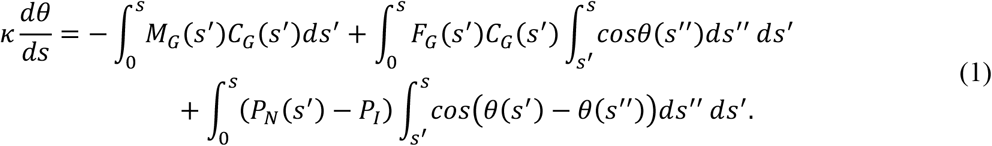

where *F*_*G*_ and *M*_*G*_ are the force and the moment respectively exerted on the membrane by individual glycoproteins, *C*_*G*_ is the local concentration of the glycoproteins, and *F*_*G*_ *C*_*G*_ and *M*_*G*_ *C*_*G*_ are the force and moment distributions respectively exerted by the glycocalyx on the membrane. Please re5fer to the next sections for treatments of *F*_*G*_, *M*_*G*_, *C*_*G*_, and *P*_*N*_. *s′* and *s*″ are dummy variables for the arc-length coordinate *ss*. Differentiation of Eq. 1 with *s* yields a more tractable force balance:

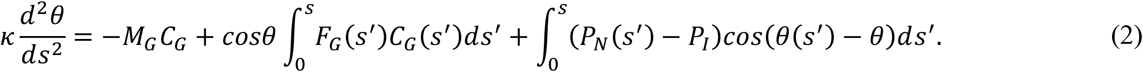

We seek solutions in the form of periodic standing waves. Thus, the solutions would be symmetric about the axis *s = L*/2. Hence, we consider a simulation domain of *s ∈* [0, *L*/2]. Periodicity at *s =* 0 and symmetry at *s = L*/2 yield zero slope boundary conditions, *θ*(*s =* 0) *= θ*(*s = L*/2) = 0. Along with these boundary conditions, Eq. 2 yields a solution for the membrane inclination.

An overall vertical force balance on the membrane equates the cytosolic force with the reaction forces from the glycocalyx and the glycocalyx counterion osmotic pressure:

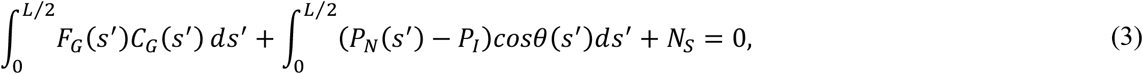

where *N*_*S*_ is a reaction force that is nonzero only at positions where the membrane touches the substrate, i.e., *H*(*s*) = 0 and prevents the membrane from penetrating the substrate. Eq. 3 indirectly provides information on the mean deformation of the membrane, which is related to the reaction forces from the glycocalyx. We configure the solver to use Eq. 3 to obtain the minimum height, *H*_0_, of the membrane. The local membrane height is then given by:

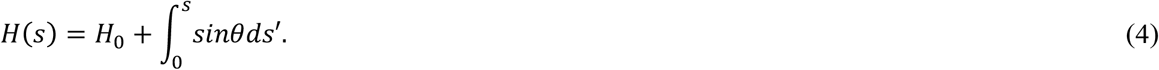

While the model formulation in this paper mainly uses a curvilinear coordinate system going across the membrane contour length, converting to a horizontal coordinate is possible using:

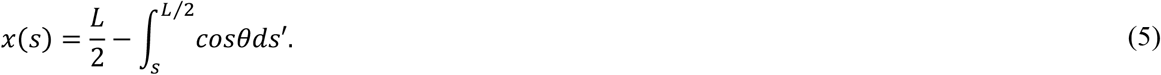

### Mechanics of beam-like glycocalyx polyelectrolytes

Glycocalyx polyelectrolytes, such as mucins, which are heavily decorated with glycan side chains, can have large persistence lengths, equal to or greater than the molecular length. In this regime, we consider the glycocalyx as a continuum of laterally mobile structural beams that bend under load. We refer to the glycocalyx constituents in this regime with interchangeable names such as “beam glycoprotein”, “beam-like polyelectrolyte”, and “bendable glycopolymer”. In this limiting case, we consider the glycocalyx to include only the beam-like biopolymers. The treatment of the glycopolymer beams is almost identical to the framework established in reference (23), with the only new element being a theoretical solution for the asymptotic limit of small beam force and small membrane inclination. We assume the transmembrane portion of a beam-like glycoprotein maintains a perpendicular orientation with the membrane due to moments from the lipids in the bilayer and interactions between polyelectrolyte side chains and the membrane. We ignore friction between glycoproteins and the extracellular substrate because observations that mucin diffusion coefficients are comparable to those of smaller transmembrane molecules (2). Thus, we consider the reaction forces to be purely normal to the substrate. Electrostatic interactions between charged glycoproteins are screened by the counterions in the interstitial fluid. With these approximations, we apply non-linear beam bending theory from fundamental solid mechanics to describe the shape of a deformed beam-like glycocalyx polyelectrolyte due to external compressive stresses (28–30):

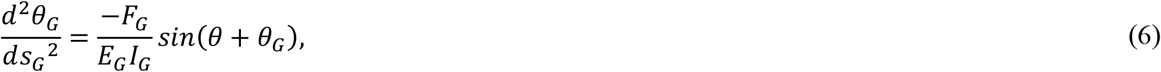

where *θ*_*G*_ (*s*_*G*_) is the protein inclination with respect to the local normal to the membrane at an arc-length coordinate *s*_*G*_, *E*_*G*_ *I*_*G*_ is the bending modulus of the glycoprotein, *F*_*G*_ is the force on the glycoprotein and *θ* is the membrane inclination at the location where the glycoprotein is attached to the membrane. Fig. 2B demonstrates a generic configuration of the beam-like glycopolymers embedded in the membrane. As described previously, we assume internal force dipoles in the lipid membrane to be enough to keep the attachment of the beam glycopolymers with the membrane oriented perpendicular to the membrane.

In the absence of other force dipoles, the boundary condition at the extracellular end is *dθ*_*G*_/*ds*_*G*_ = 0 at *s*_*G*_ *= L*_*G*_, where *L*_*G*_ is the length of the glycoprotein. The glycoprotein is constrained to be perpendicularly attached to the membrane through the boundary condition *θ*_*G*_ (0) = 0. A solution of Eq. 6 is:

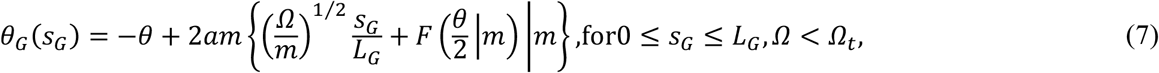

where 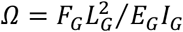 is a dimensionless force on a glycoprotein, *am*(*w* ∨ *m*) is the Jacobi amplitude function with argument *w* and parameter *a*, and *F*(*ϕ*|*m*) is the incomplete elliptic integral of the first kind with argument *ϕ. m =* (*sin*((*θ* + *θ*_*G*1_)/2))^−2^, where *θ*_*G*1_ *= θ*_*G*_(*s*_*G*_ *= L*_*G*_) is the inclination of the extracellular end of the beam. *Ω*_*t*_ is a dimensionless transition force described below. Note the slight correction in the formula for *a* relative to the one in reference (23) to adjust for a missing exponent.

For forces below a transition value *Ω*_*t*_, *θ*_*G*1_ and *a* can be determined by solving the equation *θ*_*G*_ (*s*_*G*_ *= L*_*G*_) *= θ*_*G*1_. *Ω*_*t*_ is given by:

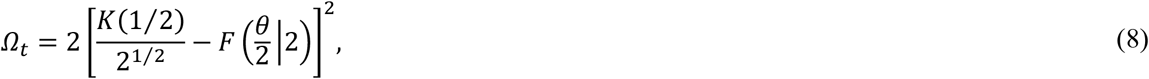

where *K*(*m*) *= F*(*π*/2 |*m*) is the complete elliptic integral of the first kind. There is a minor correction in Eq. 8 relative to reference (23), taking care of a missing exponent.

Beyond the transition force, i.e. for *Ω*≥*Ω*_*t*_, a part of the beam aligns with the extracellular substrate starting from its extracellular end, as illustrated in Fig. 2C. Note that the glycoprotein is still connected perpendicularly to the membrane but is merely aligned with the extracellular substrate due to the large force acting on the beam. In this case, *m =* 2 for the part of the glycoprotein that can deform, i.e. 0≤*s*_*G*_ ≤*s*_*GA*_, where *s*_*GA*_ is the arc length coordinate beyond which the glycoprotein is aligned with the substrate. The aligned part of the glycoprotein does not deform and thus, *θ*_*G*_ *= π*/2 − *θ* for *s*_*GA*_ ≤ *s*_*G*_≤1. The solution in Eq. 7 yields glycoprotein shapes equivalent to those obtained by a finite difference solution of Eq. 6 and with much less computational cost.

For accurate descriptions of the mechanics of the glycocalyx beams at small deformations, we use a simplified equation in the case where the deflection of the beam is small (*cosθ* − *H*/*L*_*G*_ *≪* 1) and the inclination of the membrane is also small (*θ ≪* 1). In this case, the dimensionless reaction force, *Ω*, on the glycocalyx beam can be calculated for a given glycocalyx height by using:

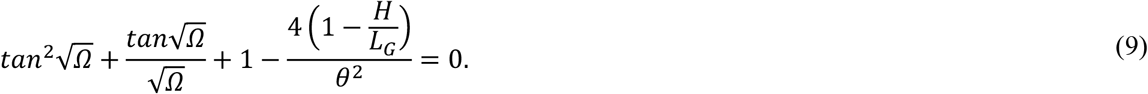

### Steric interactions and entropic pressures for beam-like glycopolymers

The components of the glycocalyx, including various mucins, are bottlebrushes with side chains attached to a backbone. Stiffer glycocalyx bottlebrushes resemble cylinders. Thus, as a first approximation, we consider these stiff beam-like elements to experience hard-disk interactions. These interactions push the beams from regions of high to regions with low glycoprotein concentrations. A virial expansion provides the osmotic pressure at low planar densities. The following Padé approximation provides an osmotic pressure that diverges as the density approaches the hexagonal closed packing limit (31):

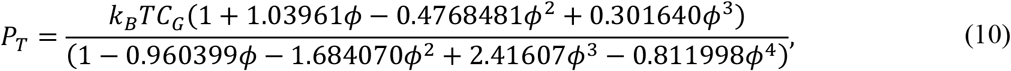

where *ϕ* is the fraction of membrane area covered by the backbones and branches of the glycocalyx polyelectrolytes. Considering each polyelectrolyte to resemble a cylindrical beam with diameter *d, ϕ = C*_*G*_ *π d*^2^/4. The effect of this tangential steric pressure on the spatial distribution of the glycocalyx is described in a subsequent section.

In a process called Donnan equilibrium, counterions from the interstitial fluid are attracted to the charged glycocalyx, resulting in an osmotic pressure of the counterions. We consider the local counterion density to be proportional to the density of the beam elements. Then the osmotic pressure of the counterions acting on the membrane normal to the surface is (32):

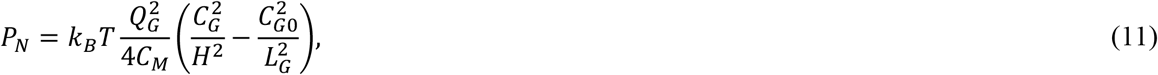

where *Q*_*G*_ is the number of counterions per polyelectrolyte and *C*_*M*_ is the ionic strength of the surrounding media.

### Polymer brush of flexible-chain-like glycocalyx polyelectrolytes

Glycocalyx polyelectrolytes with a low density of side chains can have persistence lengths much smaller than the molecular length and can behave as flexible chains. In this regime, we model the glycocalyx as a brush of flexible polymers (Fig. 2D) (33,34). Note that in this limiting case, we consider only the flexible brush glycocalyx and do not include beam-like glycoproteins. The polymers have a Kuhn length *L*_*K*_, which is twice the persistence length, an indicator of molecular stiffness. *N*_*K*_ such Kuhn segments of length *L*_*K*_ constitute a polymer. The polymers are allowed to migrate on the plane of the membrane due to compressive forces and diffusion.

The free energy of a compressed polymer brush contains translational and configurational entropies of the polymers as well as energetic contributions from excluded volume and counterion effects. The total free energy per area of the brush is (32,35):

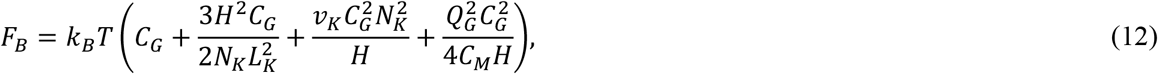

where *𝒱*_*K*_ is the excluded volume of a Kuhn segment. Given the bottlebrush nature of many elements of the glycocalyx, we consider cylindrical segments with diameter κ and length *L*_*K*_. Then the excluded volume of each cylindrical segment is given by 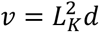 (36), which can be derived using excluded volume theory. Note that the resulting expression for cylinder excluded volume differs from the formula for the cylinder volume. Appropriate derivatives of the free energy density in Eq. 12 provide expressions for the pressures experienced by the polymers and the membrane. An osmotic pressure in the plane of the membrane that causes migration of the chains is given by,

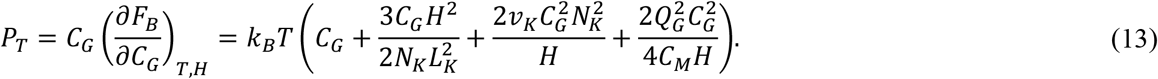

This tangential 2D pressure can impact the spatial organization of the glycocalyx as discussed in the next section. On the other hand, the membrane experiences a reaction pressure normal to the surface due to the compression of the brush,

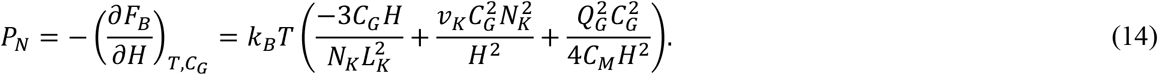

Eq. 13 and Eq. 14 provide polymer brush reaction forces due to compression. *F*_*G*_ = 0 and *M*_*G*_ = 0 for the polymer brush. In the absence of external compression, i.e. *P*_*I*_ = 0, the brush is at equilibrium when the normal pressure driving extension or compression is zero, *P*_*N*_ = 0. Considering a uniformly distributed brush, *C*_*G*_ *= C*_*G*0_, the equation *P*_*N*_ = 0 yields the stress-free height of the glycocalyx brush:

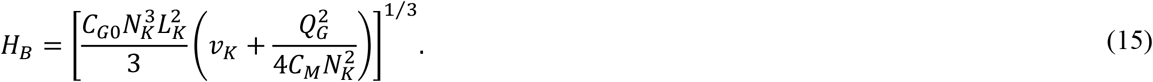

*H*_*B*_ serves as a length scale for characterizing the height of the brush glycocalyx in the presence of external compression.

This theoretical treatment considers the glycocalyx as a brush of flexible chains alongside the shape equation describing the membrane deformation. However, the brush free energy (Eq. 12) does not consider effects of large membrane curvature and is valid mainly for small membrane bending. Large membrane curvatures would require the incorporation of lateral spatial heterogeneities on the scale of the brush height leading to non-local interactions between the flexible chains. Including these effects would require a description of polymer brush behavior on a surface with spatially varying curvature, which is not currently available.

### Dynamics of glycocalyx transport

The compressive stress on the glycocalyx due to cytosolic pressure can induce migration of the polyelectrolytes in the glycocalyx. To capture the resulting spatial arrangement of the glycocalyx, we consider a continuum of identical glycocalyx molecules able to migrate laterally on the membrane. The diffusion and pressure-driven migration of the glycocalyx polyelectrolytes are the rate-limiting processes in the compression of the glycocalyx. Hence, an unsteady transport equation can capture the dynamic spatial distribution of the glycocalyx polyelectrolytes:

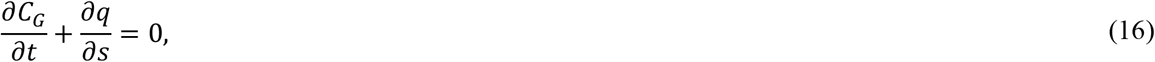

where *q*(*s*) is the local flux of the glycocalyx species:

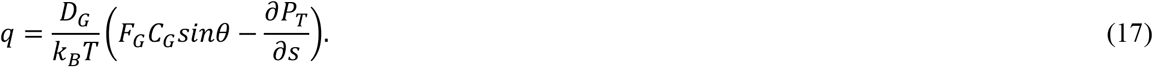

The transport equation thus becomes:

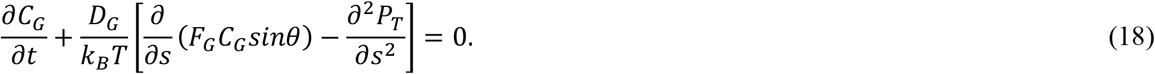

Zero flux boundary conditions are required to maintain a constant total content of glycocalyx constituents in the domain. The zero flux boundary conditions are also apparent due to the symmetry of the system around *s =* 0. Hence, the concentration profile is constrained by:

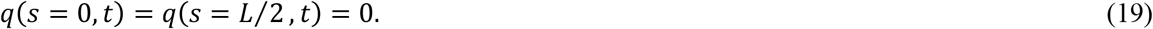

We consider a uniform concentration profile at the initial state:

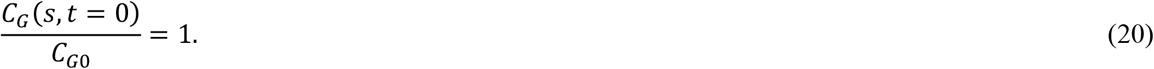

### Gibbs free energy of the beam glycocalyx

As shown in the next section, the simulations demonstrate multiple solutions to the coupled nonlinear differential equations detailed above. We determine the most favorable initial state by finding the solution with the lowest Gibbs free energy. At the initial state, we consider the glycocalyx to be uniformly distributed. Thus, the Gibbs free energy at *t =* 0 includes the bending energies of the membrane and the glycocalyx beams and the work done by the cytosolic pressure. The Gibbs free energy per membrane area with a reference state of an undeformed and uniformly distributed glycocalyx is:

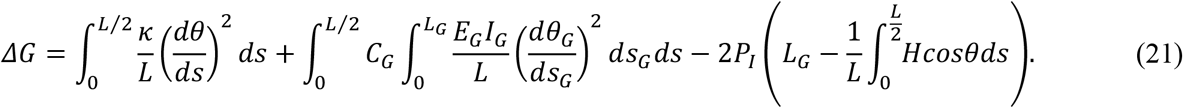

### Dynamic simulation procedure

Coupling the modeling components above allows simulation of the dynamics of the glycocalyx under cytosolic compression. We apply an approach similar to the finite volume method in order to conserve the total content of glycocalyx constituents while solving the transient transport equation (Eq. 18). Starting from the initial condition in Eq. 20, we use a fourth order Runge-Kutta solver to integrate Eq. 18 in time and calculate the dynamic glycocalyx distribution (Fig. 3). At each time step, Eq. 1-4 solve for the membrane topography. The membrane shape and glycocalyx density profile allow calculation of the normal and tangential forces exerted by the glycocalyx (Eq. 7, 9-11, 13, 14). Table 1 lists physiologically relevant values of the parameters used in the simulation. Thus, we simulate the time-dependence of the glycocalyx concentration distribution, membrane topography, and the glycocalyx forces in the presence of cytosolic pressure.

### Glossary of parameters

**Table 1.** A glossary of some of the parameters in the model, alongside physiologically relevant estimates for the same obtained from the listed references.

| Parameter name | Description | Estimated range | References |
| --- | --- | --- | --- |
| $P_I$ | Cytosolic stress on the glycocalyx | 75-450 Pa | (21) |
| $L$ | Membrane contour length scale | 200-300 nm | (37,38) |
| $C_{G0}$ | Spatially averaged glycocalyx number density | 1000 molecules/$\mu\text{m}^2$ | (9) |
| $D_G$ | Diffusivity of glycocalyx molecules on the membrane | 0.2 $\mu\text{m}^2/\text{s}$ | (2) |
| $\kappa$ | Membrane bending modulus | 200 pN.nm | (39-42) |
| $E_G I_G$ | Individual beam bending modulus | 100 pN.nm<sup>2</sup> | (29,43-48) |
| $L_G$ | Beam polyelectrolyte length | 200 nm | (49-51) |
| $d$ | Representative diameter of a glycocalyx molecule | 15 nm | (52) |
| $Q_G$ | Number of charged residues per beam or polymer | 200 | (53,54) |
| $C_M$ | Ionic strength of surrounding media | 150 mM | (55) |
| $L_K$ | Kuhn length of a polymer in the glycocalyx | 15 nm | (52) |
| $N_K$<br>$= L_G/L_K$ | Number of Kuhn segments in a polymer | 13 | - |

**Fig. 3.**
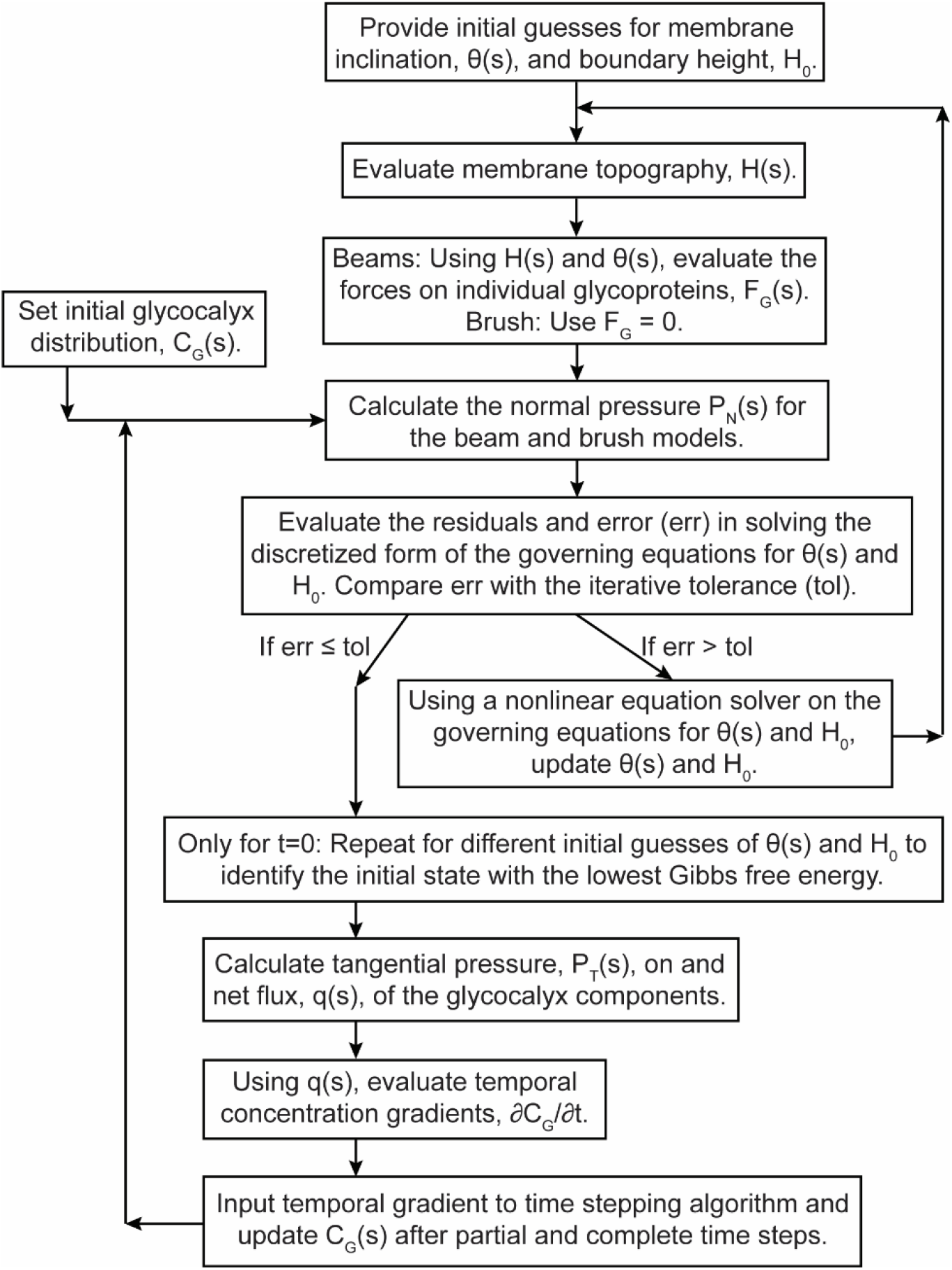
Dynamic simulation procedure. This flowchart demonstrates the time stepping process used to update the transient glycocalyx distribution and the iterative procedure used to solve for the membrane topography at each time step.

## RESULTS

### Instantaneous response of the beam glycocalyx to cytosolic pressure

We simulate the beam glycocalyx to predict the structure and organization of the glycocalyx under cytosolic pressure. At *t =* 0, the algorithm starts with user-specified initial guesses for the membrane topography. The membrane is free to translate vertically and bend, but it is constrained to have zero slope at the boundaries. The model system responds instantaneously to the application of the cytosolic pressure. To capture this response, we search for the membrane shape at *t =* 0 using a free energy minimization. The most favorable initial state corresponds to the configuration the model system would adopt instantaneously when a cytosolic pressure is applied. As described below, the interplay of physical processes results in immediate curvature of the cell membrane and deformation of the glycocalyx.

Minimizing the Gibbs free energy per membrane area for varying membrane contour length (Fig. 4A) shows the emergence of a wave-like deformation of the membrane occurring at characteristic length scales around 200-300 nm for physiologically relevant cytosolic pressures. The free energy density exhibits a minimum as a function of the membrane contour length (Fig. 4A). Thus, there exists a favored length, *L = L*_*D*_, for the membrane to deform at for every cytosolic pressure (Fig. 4A, S1). *L*_*D*_ is the contour length of the membrane domain that is the most favorable length scale for membrane bending at *t* = 0 and correspondingly, 1/*L*_*D*_serves as a characteristic scale for initial membrane curvature. The characteristic length scale of deformation decreases, and the membrane curvature increases with elevation in cytosolic pressure (Fig. 4B). Considering a dimensionless cytosolic pressure 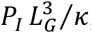, a scaling law of 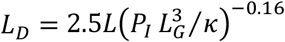. describes the trend accurately. While there is no apparent physical explanation for the exponent of –0.16 or the coefficient of 2.5, the scaling law demonstrates the power-law relationship between the cytosolic pressure and the favored length scale of deformation. Note that this is merely a description of the initial response of the model system, and temporal dynamics starting from the initial state are explored in further sections of this manuscript.

**Fig. 4.**
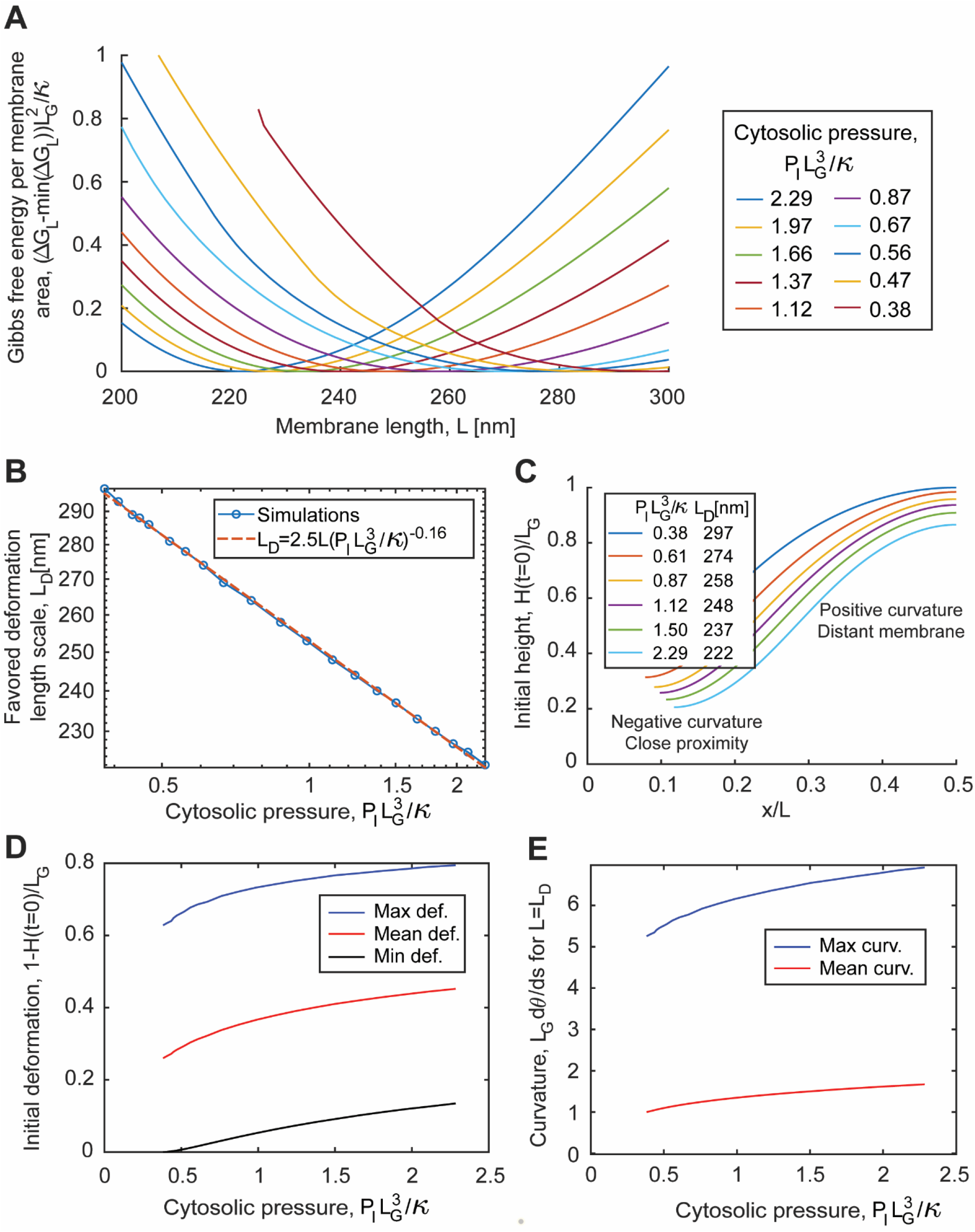
Gibbs free energy minimization at *t =* 0 reveals favorable deformation length scale for a uniformly distributed beam glycocalyx (related to Fig. S1). **A**. Plots of Gibbs free energy per membrane area against membrane contour length and cytosolic pressure provide information on favored initial states of the beam glycocalyx. The length at which the Gibbs free energy is minimum is referred to as the favored deformation length, *L*_*D*_. The Gibbs free energy curves in this figure have been translated such that the minimum value is 0 for all curves. The reference state for the free energies is a uniformly distributed glycocalyx with zero deformation. **B**. The membrane contour length at which the Gibbs free energy profile exhibits a minimum represents the favored length scale for cell surface deformation. The favored deformation length can be fit by a linear curve on a log-log plot. **C**. Initial membrane topographies for various pressures and the favored membrane deformation length. **D**. Spatially minimum, mean, and maximum deformation of the membrane at *t =* 0 show increasing trends with the cytosolic pressure. **E**. The spatially maximum and mean membrane curvatures at *t =* 0 are higher at elevated cytosolic pressures. The simulations in this figure are at *t =* 0 and use *C*_*G*0_*L*^2^ = 10, *L*_*G*_ /*L =* 1, *E*_*G*_ *I*_*G*_ /*L*κ = 1, *d*/*L =* 0.1, and *Q*_*G*_ = 200. For the parameter values used here, 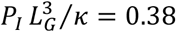 and 2.29 correspond to *P*_*I*_ = 76 Pa and 457 Pa.

The interplay between the components of the Gibbs free energy density (Eq. 21) is as follows. The internal energies of the membrane and glycocalyx compensate for the work done by the cytosolic pressure in deforming the membrane and glycocalyx. The area densities of both these contributions increase with increase in membrane contour length. The competing free energy contributions result in the existence of a minimum net free energy density at *L = L*_*D*_(Fig. S1B).

The favored deformation length, *L*_*D*_, is a contour length of the membrane domain, i.e. the length of the membrane along its curvature. *L*_*D*_does not describe the extent of bending deformations in the membrane. The magnitude of the bending deformations depends on *L*_*D*_as well as the cytosolic pressure. Fig. 4B looks at a range of cytosolic pressures and their corresponding *L*_*D*_. The membrane deformation increases monotonically with the cytosolic pressure (Fig. 4C-D). The initial membrane curvature is also larger at higher cytosolic pressures (Fig. 4E).

This section summarized the instantaneous response of the membrane and the glycocalyx when a cytosolic pressure is applied. Treatments of the dynamic changes and the equilibrium states are below.

### Dynamics of the beam glycocalyx

Starting from the instantaneously created states described in Fig. 4, the model system evolves with time through transient processes, namely the transport of the glycocalyx polyelectrolytes. The unsteady transport equation, uniform concentration profile at *t =* 0, and zero flux boundary conditions provide the dynamic glycocalyx distribution (Eq. 18-20). The membrane topography starts from the most favorable deformed state (*L = L*_*D*_) at the specified cytosolic pressure and is developed with every update of the glycocalyx concentration distribution. Fig. 5 shows the transient membrane topography and concentration profile for a sample cytosolic pressure. The compressive stresses on the glycocalyx beams in confined regions result in driving forces for convective transport of the beams to more favorable locations. This transport of the glycopolymers leaves behind emptier locations on the surface, allowing the membrane to bend further due to cytosolic pressure (Fig. 5A). The temporally increasing deformation of the membrane further pushes the regions on the membrane (Fig. 5B). At large times, the effective diffusion of the glycocalyx constituents becomes the dominant transport process. The steric interactions between glycocalyx components resist the intense concentration of the glycocalyx and, along with diffusion, drive the glycocalyx towards a uniform distribution in accessible parts of the membrane-substrate gap. The membrane, however, reaches sharply deformed states, resulting in complete exclusion of the glycopolymers from low-lying regions on the membrane. Together, these effects result in a glycocalyx distribution with two distinct phases. The transport of the mucinous fibers occurs with two characteristic time scales. For the parameters considered in this paper, the convection time scale, 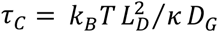, is in the range 0.01-0.02 seconds and the diffusive time scale, 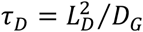, is about 0.2-0.5 seconds.

**Fig. 5.**
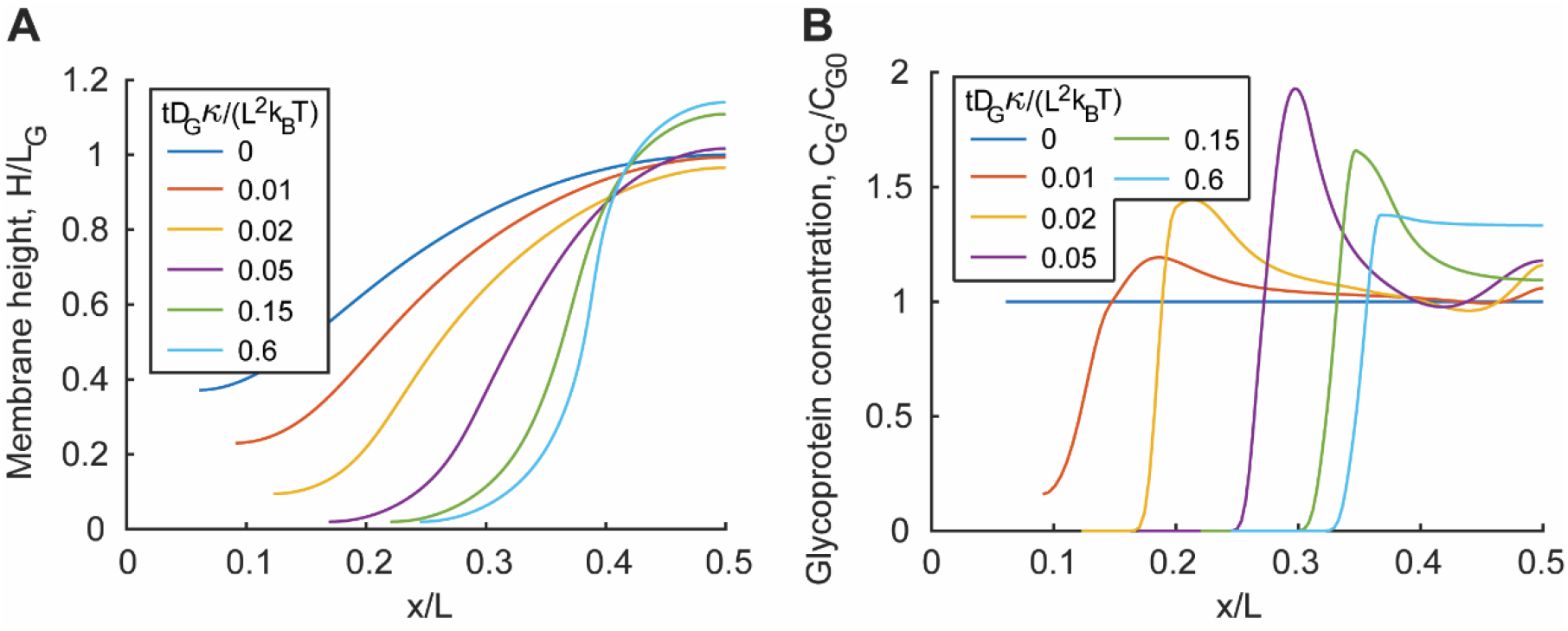
Dynamic simulations predict transient beam glycocalyx organization and membrane shape. The membrane topography **(A)** and the glycopolymer concentration profile **(B)** in Cartesian coordinates for several times in a dynamic simulation with 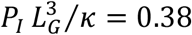 and *L = L*_*D*_= 297*nm*.

### Mechanical equilibrium of the beam glycocalyx under cytosolic pressure

The system approaches a steady state at large time as the transients die down. Fig. 6 shows the membrane topography and the beam glycocalyx distribution at equilibrium. The predicted equilibrium state shows a highly deformed membrane topography with sharp gradients (Fig. 6A). Relative to the initial membrane curvatures (Fig. 4E), the membrane curvatures at equilibrium (Fig. 6B) are heightened. The equilibrium curvatures also increase with the cytosolic pressure (Fig. 6B). At lower pressures, the membrane contains areas where the height exceeds the glycopolymer length. The glycopolymers in these elevated areas do not offer any mechanical resistance as they do not contact the substrate. In these cases, the glycocalyx polymers in regions with smaller membrane height bear the cytosolic pressure. The glycocalyx is completely pushed out from the membrane areas with low heights and exhibits a nearly uniform distribution in the rest of the membrane (Fig. 6C). Surprisingly, the concentration profiles for different pressures almost coincide when plotted against the membrane arc length coordinate (Fig. 6D).

**Fig. 6.**
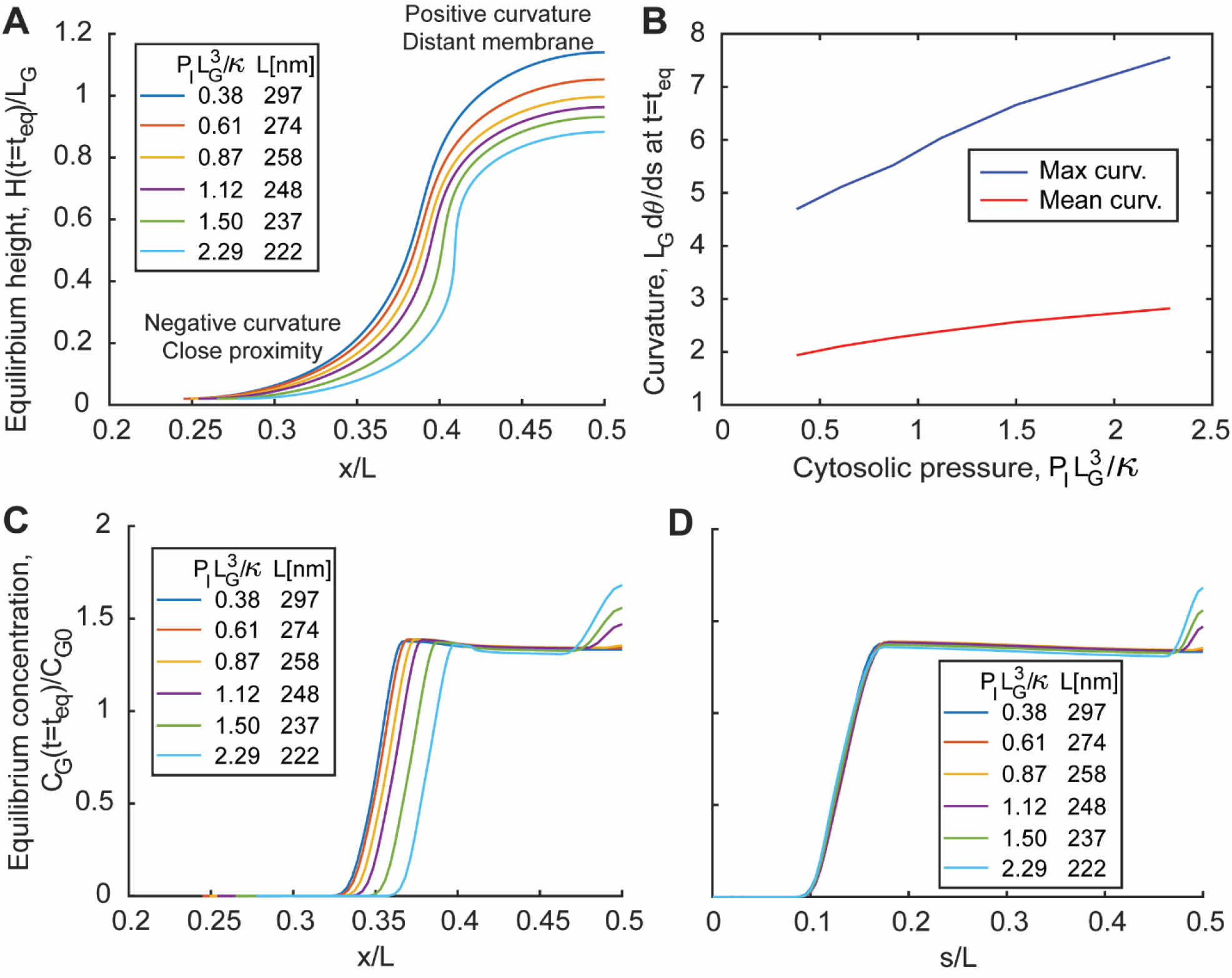
Beam glycocalyx structure and organization at steady state. **A**. Equilibrium membrane topographies in Cartesian coordinates for a range of cytosolic pressures and favored deformation lengths. **B**. The spatially maximum and mean membrane curvatures increase with cytosolic pressure and are higher at equilibrium than at the initial state. **C**. Steady state glycocalyx concentration profiles against the horizontal spatial coordinate, *x*. **D**. The steady state profiles for different cytosolic pressures plotted as a function of the membrane arc length coordinate nearly coincide.

### Mechanics of a cytosolically compressed brush glycocalyx

To capture the behavior of a glycocalyx containing flexible elements, we simulated the polymer brush glycocalyx. Starting with the same initial condition of a uniform spatial distribution of the glycocalyx polyelectrolytes, we simulate the dynamic behavior. Unlike the beam glycocalyx model, the initial state for the brush exhibits an undeformed cell membrane (Fig. 7A). The brush itself, however, is deformed and the height (*H*) of the compressed brush can be calculated by solving an algebraic equation derived by balancing *P*_*N*_ *= P*_*I*_ and considering *C*_*G*_ *= C*_*G*0_:

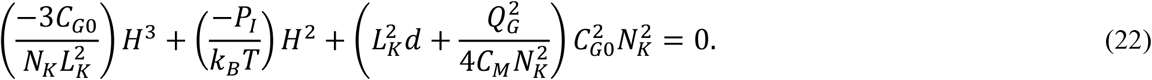

**Fig. 7.**
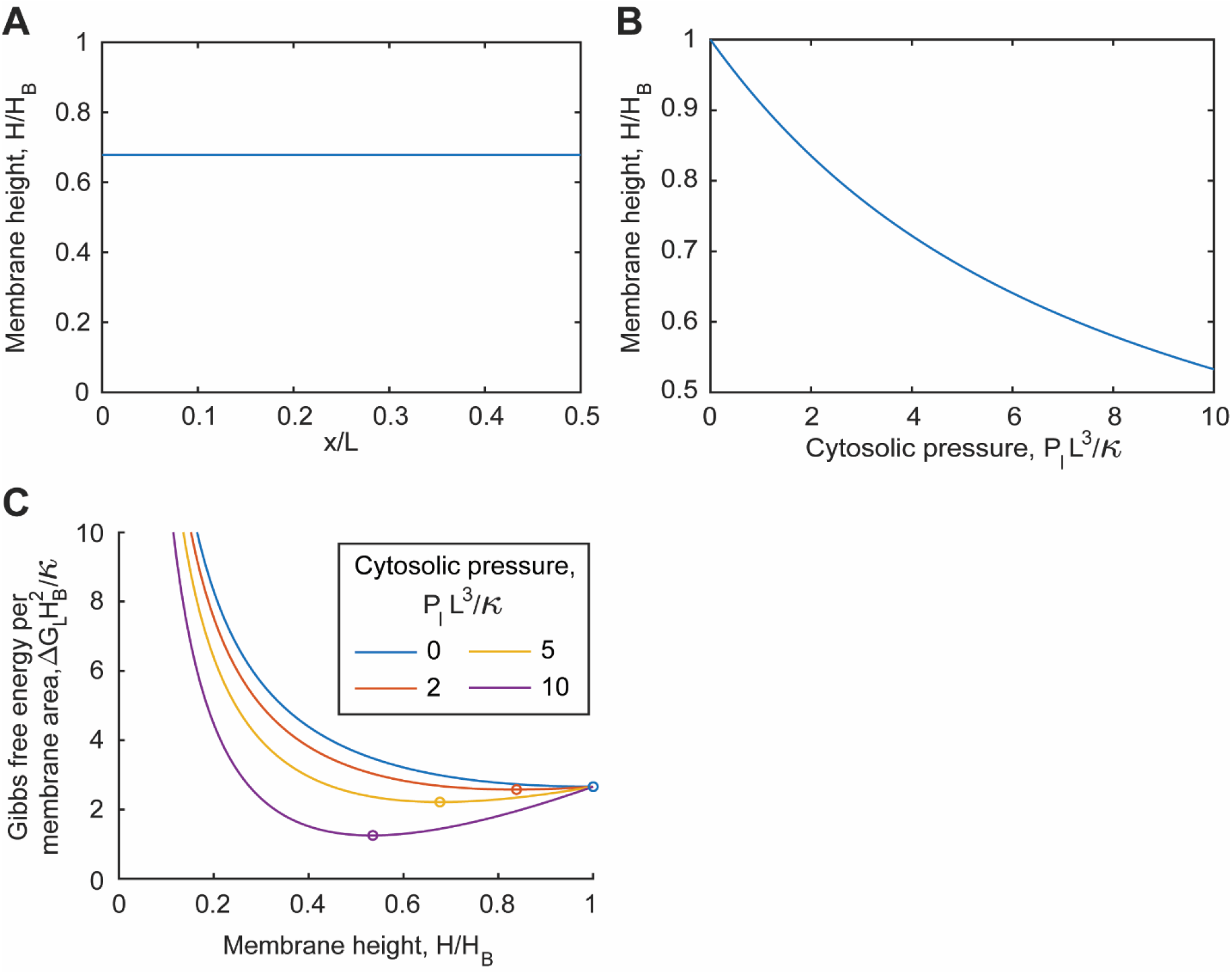
Simulations of the brush glycocalyx model. **A**. Membrane topography for an example simulation using *P*_*I*_ *L*^3^/κ = 5 and *L =* 250*nm*. **B**. Solution of Eq. 22 provides the membrane height as a function of the cytosolic pressure. **C**. Gibbs free energy of the brush glycocalyx, including the work done by the cytosolic pressure for varying membrane height exhibits minima.

The brush height decreases with increase in pressure (Fig. 7B).

The Gibbs free energy of the model system for varying membrane height helps interpret the prediction of an undeformed membrane. The Gibbs free energy for a uniformly distributed glycocalyx and a flat membrane contains the brush free energy, *F*_*B*_ (Eq. 12), and the work done by cytosolic pressure. The work done by cytosolic pressure is directly proportional to the change in volume of the glycocalyx and can be calculated by taking the difference of i) the original volume and ii) the volume at any instant, evaluated by integrating the membrane height over the x-dimension. The total Gibbs free energy is:

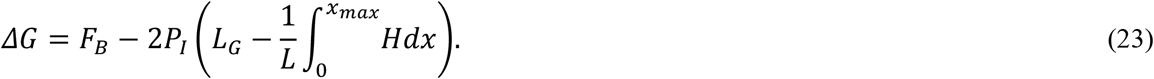

*x*_*max*_ can be calculated using:

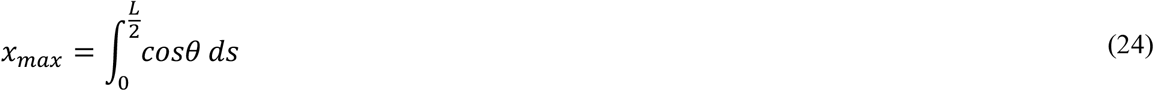

The Gibbs free energy exhibits a minimum against membrane height (Fig. 7C), suggesting the presence of an optimal height for every point in the membrane to adopt. The polymer brush calculation predicts a flat membrane with this optimal height. Any deformation to the brush from the flattened state would result in an increase in the Gibbs free energy and would be unfavorable. However, the polymer brush model in this paper ignores non-local effects and is valid only for small membrane curvatures. Incorporating more advanced effects may lead to predictions of deformed brush states for uniformly distributed glycocalyces.

There is a lack of dynamics in a cytosolically compressed brush glycocalyx. As the membrane height, *H*, and the glycocalyx polyelectrolyte concentration, *C*_*G*_, are both spatially uniform, the tangential pressure, *P*_*T*_, is also uniform. Thus, convective forces driving migration of the polyelectrolytes are absent. Diffusion is the dominant transport process and drives the glycocalyx distribution to maintain its initial uniform concentration profile.

## CONCLUSION

In this paper, we present a numerical model for the dynamic reorganization of the glycocalyx in the presence of cytosolic pressure and interacting with an external surface. The model considers the mechanics of the structural elements in the glycocalyx tethered to the cell membrane and interacting with an extracellular substrate. Two limiting modeling frameworks, bendable beams and flexible chains, are considered to cover the diverse range of stiffness of naturally present glycocalyx polyelectrolytes:. Application of a cytosolic pressure results in instantaneous deformations with characteristic membrane deformation length scales and membrane curvatures. The favored deformation lengths are inversely related to the cytosolic pressure through a power law. Transient calculations demonstrate the growth of the membrane deformation and gradual exclusion of the glycocalyx beams from areas of the membrane closest to the substrate. Steady states with sharp membrane topographies and nearly biphasic glycocalyx concentration profiles are observed. The concentration profiles suggest the formation of clusters of the glycocalyx beams. The polymer brush model predicts a uniformly compressed brush and an undeformed cell membrane. The height of the compressed brush decreases with increase in cytosolic pressure.

Blebbing motility is a fast, low-adhesion, amoeboid migration mode. In particular, blebbing motility is used effectively by tumor cells, immune cells, and embryonic cells to traverse through 3D extracellular matrices and tissues. The glycocalyx has been known to sterically block interactions between cell surface receptors and the cell environment (3). Elevated cytosolic pressures during blebbing motility must be experienced by the glycocalyx at the cell-environment interface. In this paper we demonstrate that the glycocalyx is quickly (∼0.01-0.02 s) redistributed upon the application of cytosolic pressures, leaving behind sparsely covered membrane areas, which are pushed by the cytosol to generate large membrane deformations. The membrane deformations are large enough to allow for the formation of adhesion complexes through cell surface receptors such as integrins. Blebs grow on a time scale of ∼30 s and retract back in another ∼120 s (22). These results suggest that the glycocalyx does not prohibit adhesion formation on blebs with elevated cytosolic pressure. In this paper, the glycocalyx polyelectrolytes are freely mobile with their motion resisted only by the viscosity of the cell membrane and steric interactions with other glycocalyx polymers. However, immobile transmembrane entities, such as proteins, might directly hinder and hydrodynamically influence the translation of the glycocalyx molecules. Immobile proteins on the cell membrane are known to appreciably reduce rates of diffusion of mobile transmembrane molecules (56), which may be contributing to the larger time scales observed in blebbing. Additionally, the glycocalyx elements can interact biochemically with the extracellular matrix and mediate glycan-dependent cell attachments (57). Mucins also can interact with each other through multivalent lectins called galectins (58,59). Overall, these effects would impact the rates of glycocalyx redistribution, influence the generation of large membrane deformations, and affect the propensity for cell adhesion.

The density of glycocalyx polyelectrolytes also may serve as a regulator for cell adhesion. In a prior theoretical model, we showed that increasing the glycocalyx density enhanced the mechanical resistance offered by the glycocalyx against cytoskeletal compression (23). There may be a similar effect of the glycocalyx density on the resistance against cytosolic pressure. Further investigations considering immobile glycocalyx elements and varying glycocalyx densities are necessary to address these questions.

Membrane topography plays important roles in various facets of cellular dynamics, including the signaling pathways originating from curvature-sensing molecules (10), the organization of intracellular actin (11), and the activation and binding of cell adhesion receptors and subsequent signaling (2). Of particular relevance here are curvature-sensing proteins that are recruited and activated by membrane bending. The curvature-sensing F-BAR protein, FBP17, is responsible for stimulating actin polymerization through Arp2/3 in cells butted against nanopatterns (11). FBP17 is seen to sense curvatures in the range of 1/(400 nm) to 1/(100 nm) (11). The G-protein-coupled receptor, Y2R, is recruited into highly curved membrane structures such as nerve terminals and is involved in cell migration and angiogenesis (60). Y2R has been shown to accumulate in filopodia with curvatures of 1/(100 nm) to 1/(25 nm) (60). The results presented in the current manuscript show the role of the glycocalyx in regulating the membrane shape while in the presence of a cytosolic pressure. In the case of the beam glycocalyx model, the interplay of physical processes resulted in equilibrium membrane states with mean curvatures of approximately 1/(50 nm) and maximum curvatures of about 1/(20 nm). Given the relevant magnitudes of the curvatures generated, the glycocalyx under cytosolic stresses may have implications on membrane-curvature-sensing protein activity.

We also predict the formation of periodic patterns at the cell surface with characteristic length scales (*L*_*D*_) of 200-300 nm. These periodic topographies could function as glycocalyx-derived nanopatterns for curvature-sensing molecules. Steric interactions between polyelectrolytes in dense glycocalyx brushes drive the formation of tube-like and pearl-like shapes at equilibrium (9). However, the transient mechanisms resulting in these shapes are unclear. The membrane deformations illustrated in this paper, generated by the interplay between the glycocalyx and the cytosolic pressure, may provide the initial push needed to start growing longer membrane protrusions.

The modeling formulation and its predictions presented in this paper are an advancement towards a clearer understanding of the physical functions of the glycocalyx during blebbing motility. However, a comprehensive model for the glycocalyx during blebbing motility would need the addition of further modeling components. For example, experimental data could be used to prescribe the time-dependent size of and pressure in a bleb. This would allow for the modeling of the glycocalyx on a bleb with temporally varying properties. Interfacing with an extracellular substrate, the contact area between the glycocalyx and the substrate would increase as the bleb grows and decrease as the bleb retracts. Incorporating adhesion receptors into such a modeling formulation would yield insight into the contact times needed to create adhesions for the purpose of motility. The polymer brush model for the glycocalyx would also benefit from further development. The lack of membrane deformation predicted for the brush model suggests the need for the incorporation of more complex effects not addressed in this paper. The present model ignores non-local effects, whereas in a crowded glycocalyx, flexible chains attached to the membrane at an area with a lower height may be able to explore spaces available in elevated areas of the membrane. This would allow for relaxation of the requirement of small membrane curvatures in the case of the brush model.

This article presents a biomechanical model for the glycocalyx under cytosolic pressure by incorporating dominant physical effects. The mechanics of the structural elements in the glycocalyx and the cell membrane, steric effects and counterion osmotic pressure considerations in the glycocalyx regulate the biomechanics of the glycocalyx. The convection of the structural glycopolymers due to the cytosolic pressure and the diffusion of these molecules on the membrane govern the time scales of the biomechanical response. Glycocalyces with stiff, beam-like elements exhibit the existence of favored length scales for cell surface deformation that decrease with increase in cytosolic pressure. The computations predict the gradual formation of membrane topographies with sharp spatial gradients and the formation of homogeneous glycopolymer clusters in the elevated membrane areas. The pressure-dependent deformation length and the formation of homogeneous clusters due to a cytosolic pressure may provide insight into transient processes leading to the formation of curved membrane shapes and bleb-dependent cell migration.

## AUTHOR CONTRIBUTIONS

All authors contributed to conceiving the presented idea, developing the computational model, performing computations, discussing results, and writing the manuscript.

## ACKNOWLEDGEMENTS

This investigation was supported by the National Institute of General Medical Sciences 5R01GM138692 (M.J.P), National Cancer Institute U54 CA210184 (M.J.P), and National Science Foundation Grant 1752226 (M.J.P.).

