## Supplemental Figure 1 for "Glycocalyx dynamics and membrane curvature under cytosolic pressure"

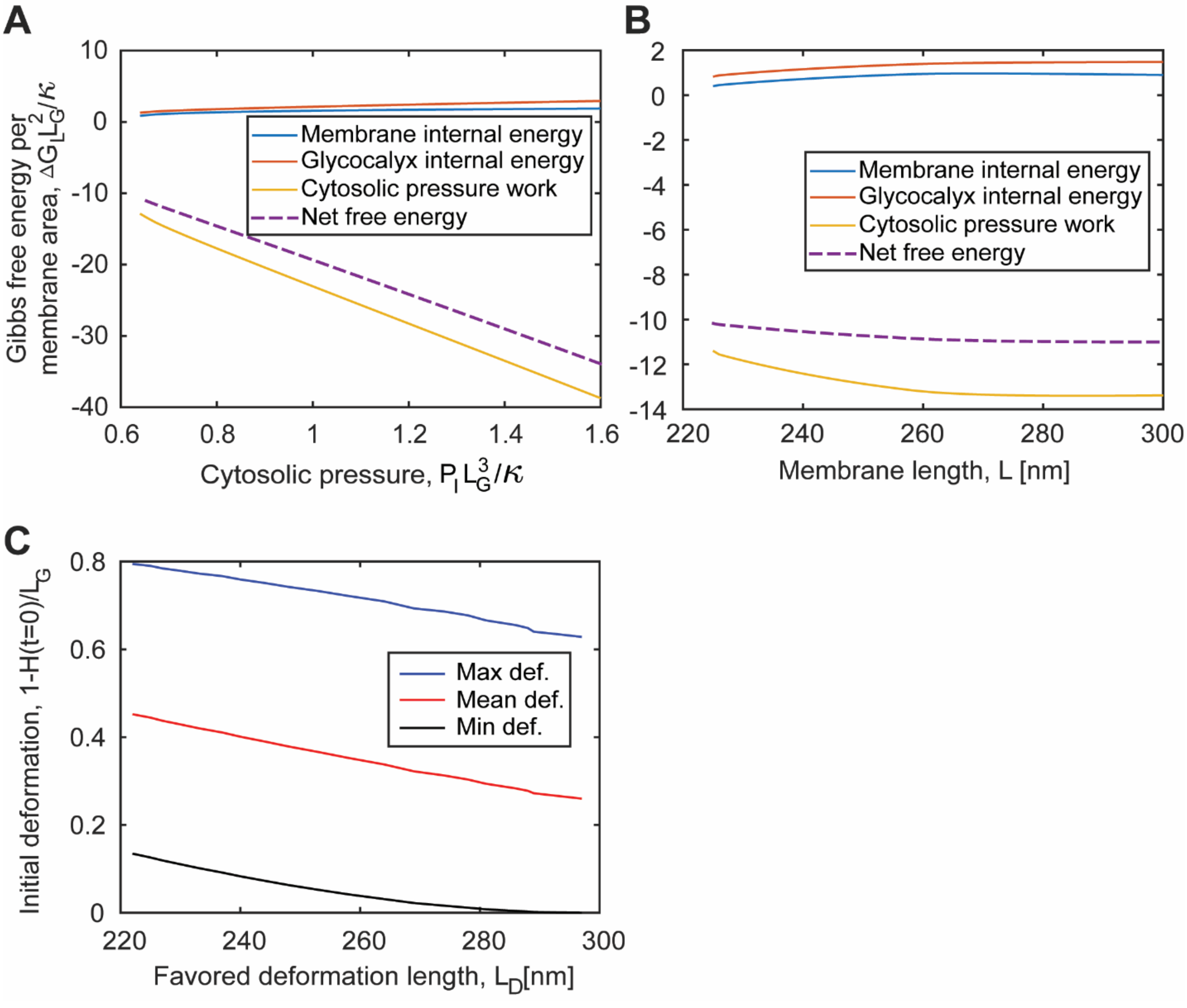


**Fig. S1 (related to Fig. 4).** Membrane and glycocalyx internal energy and cytosolic pressure work contributions to the Gibbs free energy vary with the cytosolic pressure **(A)** and the membrane domain length **(B)**. The work done by the cytosolic pressure in deforming the glycocalyx is an important contribution lowering the free energy. This work term is amplified with increasing pressure, which results in more deformed states at higher pressure **(A)**. **C.** Trends for the spatially maximum, mean, and minimum deformations against the favored deformation length at $t = 0$. These curves are generated using the information contained in **Fig. 4B** and **Fig. 4D**. Since the favored deformation length is inversely related to cytosolic pressure, the membrane deformation is smaller for deformations with larger $L_D$. This prediction is contrary to the typical positive correlation between length scale of deformation and extent of deformation of a mechanical element. $C_{G0}L^2 = 10, L_G/L = 1, E_G\, I_G/L\kappa = 1, d/L = 0.1$, and $Q_G = 200$ in this figure. $L = 250nm$ in **A** and $P_I\, L_G^3/\kappa = 0.38$ in **B**.
